# Angular Substrate Preference and Molting Behavior of the Giant River Prawn, *Macrobrachium rosenbergii* and Its Implications for Cannibalism Management

**DOI:** 10.1101/354472

**Authors:** Malcolm L. McCallum, Samad D. Weston, Yonathan Tilahun

## Abstract

The Giant River Prawn is an important commercial species from southeastern Asia and has a large global market. It has a complex life cycle in which it undergoes several molting sequences. Many arthropods require firm perches on which they can perform ecdysis. We investigated preference for substrate slope and its influence on ecdysis. We discovered that prawns occupy horizontal surfaces more frequently than others, but during pre-molt and molt stages, they shift their habitat use to non-horizontally sloped surfaces. Here, they will flex their shell and later molt. We recommend modification of cannibalism management in commercial facilities by providing sufficient vertical (strongly preferred) or high-sloped (greater than 30 degrees) surfaces to facilitate ecdysis, while providing much horizontal space for foraging and other activities. This should create habitat separation between foraging and highly susceptible freshly-molted prawns, thus leading to reduced cannibalism-related mortality.

**Funding:** This work was supported by the USDA Evans Allen Program at Langston University, Project Number USDA-NIFA-OKLUMCCALLUM2017.

**Disclosure statement:** We acknowledge that there is no financial interest or benefit that has arisen from the direct applications of our research.

## Introduction

The Giant River Prawn (*Macrobrachium rosenbergii* [De Man 1879]) is a large crustacean indigenous to southern and southeastern Asia, Oceania, and several islands in the South Pacific Ocean (New 2002). A global market for this species has developed since the 1990s, so it is commercially raised for food around the globe (FAO 2002). The species undergoes a complex life cycle involving spawning in brackish waters, hatching as planktonic larvae, and development into post-larvae followed by metamorphosis into juvenile prawns that migrate back into freshwater to mature and return to estuaries to spawn. Giant River Prawns feed on an assortment of live, dead and decaying plant and animal matter. They become aggressively cannibalistic in captivity, providing complications for producers.

Despite their widespread commercial appeal, details about their life history and behavior are not well detailed. This lack of information is important because these kinds of details form the foundation for appropriate husbandry practices and because life history details provide the outcomes of the evolutionary process, suggesting much about an organism’s biology (McCallum and McCallum 2006; Bury 2006). One of these life history elements is their aggressive and cannibalistic nature. Producers confront this management problem by creating infrastructure in ponds and aquaria to provide more habitable surface area. This is thought to dilute prawns, which limits encounters leading to improved growth rates and survivorship. Details vary about how this infrastructure should be constructed. Some producers place vertical mesh walls with enough separation to allow space for the size of prawn housed. Another common option is to create a series of horizontal mesh platforms stacked one over the other. Variations on these themes run the gamut, with many placing lines of snow fencing through ponds or bundling plastic mesh for placement in aquaria or ponds.

We asked if prawns use vertical, horizontal and angled surfaces similarly. The rationale is that investment in excess infrastructure that is not readily occupied is money and time spent unwisely on materials and maintenance. Likewise, if one uses only horizontal or vertical platforms, it may lower production potential if prawns have a biologically-based preference. We hypothesized that prawns may demonstrate preference for the angularity of the substrate. We predicted that if they do, they would be found more frequently on substrates of some angles than others.

## Methods

Between 75-100 juvenile prawns (body mass range = 0.01 − 0.045 g) were housed communally in an aerated 40-L glass aquarium containing 30-L of medium toxicological hard water (U.S. EPA 2002). Prawns were fed ground shrimp pellets daily *ad libitum*. Four 6.35 × 30 cm long strips of plastic mesh (3 mm mesh size) were formed into 30-60-90 triangles to serve as perching structures for prawns. Two were placed with the short leg and two with the long leg as the horizontal base of the triangle. Each triangle was observed twice daily (N = 9 observation periods)and ^~^7 hrs apart, for five minutes and the number of prawns observed on each surface was noted. These data were converted to prawns/mm and then angle preference was analyzed using one-way ANOVA with a Tukey means comparison test. We also recorded the number of tail flexing prawns (Fig. 1) and the number that hung upside down versus right side up on each side. Pre-molt flexing behavior was analyzed using Chi Square.

We also housed 30 prawns in individual 50 ml flasks from an unrelated experiment. Seven of these were provided vertical perches made out of 3 mm^2^ plastic mesh. Molting success, frequency and mortality were recorded over a four-week period. The data were analyzed using Chi square with an alpha = 0.05.

## Results

Data for preferred perch angle was not normally distributed (Anderson-Darling: A^2^ = 1.494, P = 0.001), so we transformed it using the “normal scores” function in MiniTab 13.0 to allow analysis by ANOVA. Although prawns used some surfaces more than others (Fig. 2; ANOVA: F_(3, 108)_ = 5.60, P = 0.002), there was no significant difference between use of 60 degree and 30 degree sloped surfaces (Tukey: −1.003, 0.3710). Thus, we pooled data to prawns resting on the hypotenuse, vertical or horizontal mesh surfaces to simplify further analyses. Here, prawns used some surfaces more than others (ANOVA: F_(2, 105)_ = 5.60, P = 0.005). They used horizontal surfaces more frequently than vertical surfaces (Tukey: −1.23, −0.196). There was no difference in use between vertical and angled surfaces (Tukey: −0.747, 0.290). They occurred on angled surfaces marginally more often than vertical surfaces (Tukey: −0.032, 1.0051). Prawns seldom hung upside down (6.4%, 25/390 observations), preferring to remain upright.

**Figure 1.**
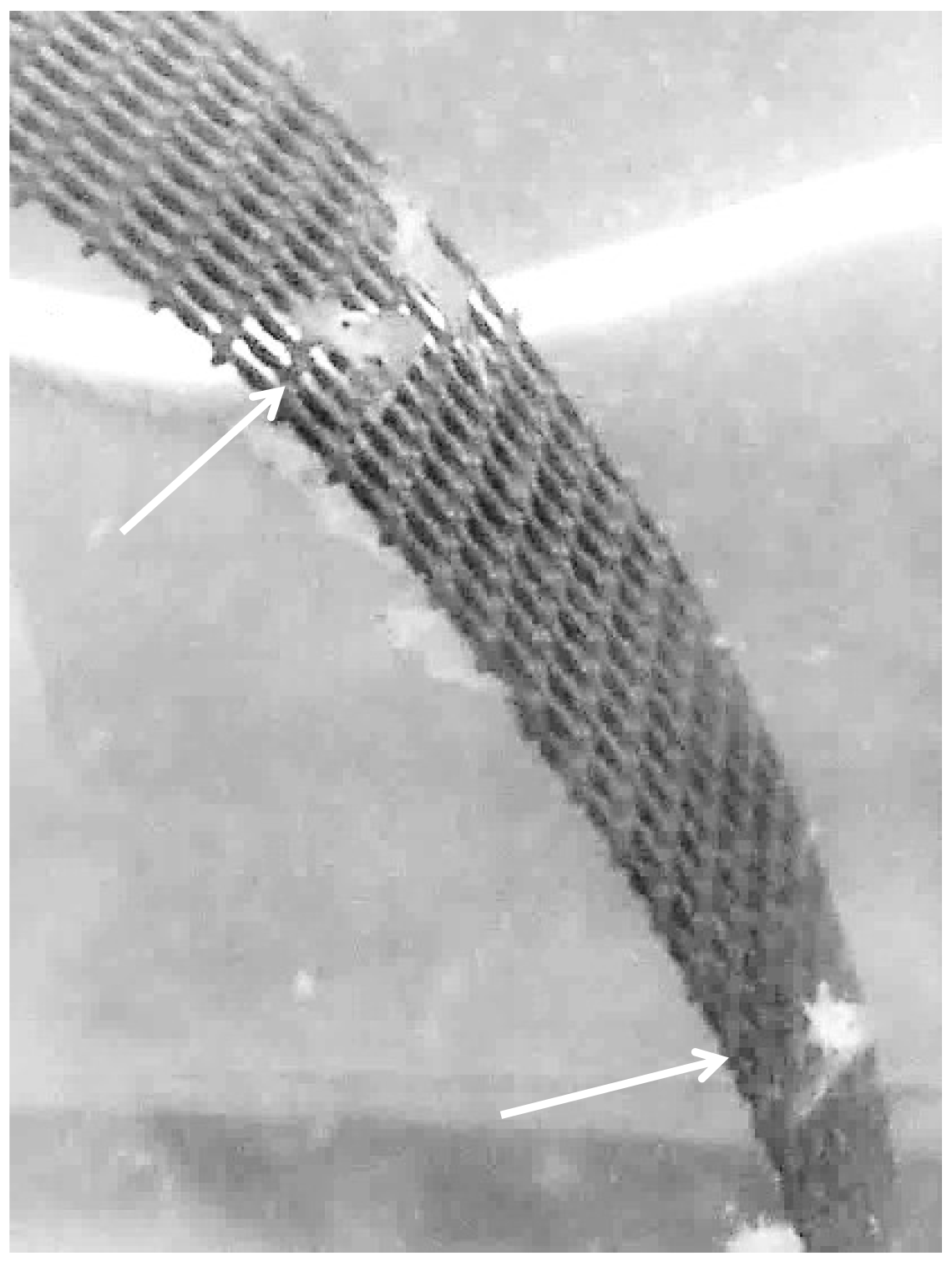
Flexing behavior of juvenile prawns on a slanted surface. Two prawns are flexing (see arrows). The third was flexing prior to taking the photograph. There are also two prawns hanging upside down.

Flexing behavior was not uniformly distributed among surfaces (Fig. 2; χ^2^ = 7.14, df = 2, P = 0.028). It occurred more frequently on vertical (χ^2^ = 7.14, df = 1, P = 0.0.008) and marginally more frequently on angled surfaces (χ^2^ = 2.78, df = 1, P = 0.096) than on horizontal surfaces. There was no significant difference in flexing behavior between vertical and angled surfaces (χ^2^ = 1.32, df = 1, P = 0.251). No prawns were observed flexing while hanging upside down.

Of the individually-housed prawns, 7 (100%) molted at least once when a perch was present. Among those not provided perches, 13/22 (59%) also molted, but the remainder failed to molt. Prawns that were not provided a perch molted less frequently than prawns that were provided a perch (Chi square = 4.15, P = 0.042).

**Figure 2.**
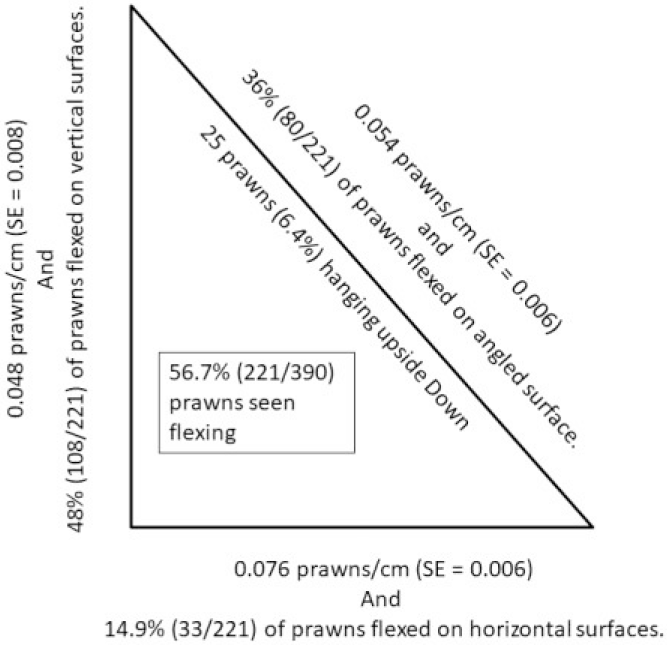
Distribution of juvenile prawns and flexing behavior on mesh infrastructure.

## Discussion

Mortality from post-larvae to adult in production systems ranges from 20 - 50% and appears related to molt state (Peebles 1978). Uniformly-sized prawns are especially susceptible to aggression and cannibalism during late pre-molt and early post-molt (Peebles 1978). Further, animals weakened by disease or environmental conditions succumb during ecdysis (Justo et al. 1991) Flexing behavior is known to precede molting in tailed decapods (Travis 1954; Tamm and Cobb 1978). Many molting insects must grasp a substrate firmly during ecdysis (Howard 1995; Fahrbach and Mesce 2005; White and Ewer 2014), much captive mortality arises when individuals in molt fall from their perch (Whitman 1986). Perhaps nowhere is this more familiar than with cicadas (Cicadidae) (Truman JW, III 2012; Truman 1983; Mantel 1971). Our results suggest a similar need in *M. rosenbergii*, and possibly other tailed decapods. Generally, the focus for captive prawn mortality has been aggression and cannibalism; however, our results beg to question if cannibalism is a symptom of lacking appropriate habitat for molting. Molting prawns occupy habitat used less frequently by non-molting individuals and molting frequency is inhibited when perches are lacking, but no increase in survivorship occurs when a perch was present. This suggests cannibalism results from the lack of non-horizontal perches that force molting prawns to occupy habitat that non-molting prawns frequent; leading to higher exposure to foraging prawns that can cannibalize newly molted individuals. However, a previous study in which surface area was increased 20% by installation of PVC frames with horizontal plastic mesh and vertical suspended seines did not significantly change mortality rates, but ponds without infrastructure had a higher proportion of small males, lower proportion of orange-clawed males, and larger body size in blue and orange clawed males, and reproductive and virgin females (Tidwell et al. 2007). Our data support these findings and may provide evidence that increasing the surface area more than 20% may provide even better results. All producers and others raising prawns in captivity should ensure that both horizontal and non-horizontal surfaces (preferably vertical, since these are used most during molting and least during foraging) are available in culture chambers to reduce molt-related mortality due to cannibalism.

## Acknowledgements

Many thanks to Steve Hart, Arthur Goetsch, Ali Hussein, Tilahun Sahlu, and Walter Meshaka, Jr. for feedback and discussions about this research. The research was funded through the USDA Evans Allen Program, Project Number USDA-NIFA-OKLUMCCALLUM2017 and was conducted under accordance with the Federal Animal Welfare Act of 1966 (Amended in 1970), Langston University IACUC #2018-11.

